# Environmental gradients and habitat specificity structure benthic microbial assemblages in a temperate seagrass ecosystem

**DOI:** 10.64898/2026.01.05.697649

**Authors:** Gabriel A. Serrano, Alejandro De Santiago, Tiago José Pereira, Mirayana Marcellino-Barros, Holly M. Bik

## Abstract

Seagrass meadows are globally distributed coastal ecosystems that provide habitats for diverse species and facilitate ecosystem services, such as carbon storage and nutrient cycling. However, factors linked to human activities and climate change are causing seagrass decline worldwide, thus impacting benthic community structure and ecosystem functioning. Although macrofaunal communities in seagrass habitats are well studied, the structure and function of bacteria/archaea and microeukaryote assemblages remains underexplored. In this study, we used environmental DNA (eDNA) metabarcoding (16S and 18S rRNA genes) to analyze bacteria/archaea (16S), microeukaryotes (18S raw sediment), and meiofauna (18S Ludox) communities in seagrass and bare sediment habitats across three sites along an estuarine gradient in Bodega Harbor, California. We found that microbial and microeukaryote communities were primarily structured by site as a response to the environmental gradient (e.g., sediment properties), whereas meiofauna responded more strongly to the presence of seagrass. Beta diversity showed strong differentiation among estuarine sites, while alpha diversity varied within sites, with bare sediments often displaying higher diversity. Network analysis revealed significant co-occurrences between bacteria and nematodes, including chemosynthetic symbiont-bearing lineages in mudflats and stress-tolerant bioindicators. Our findings underscore the importance of environmental gradients and microhabitat features in shaping benthic biodiversity in temperate seagrass ecosystems. Furthermore, it highlights potential microbial-nematode associations, likely a response to environmental variation. By integrating multiple benthic components, our study advances understanding of biodiversity drivers in temperate estuarine seagrass habitats and provides new insights on microbial community structure in these ecosystems.

**Importance:** Seagrass meadows, which sustain a wide variety of plant and animal life, are vital coastal habitats. These submerged ecosystems support biodiversity, protection of coasts, nitrogen cycling, and carbon storage. However, they are declining due to pressures from climate change and human activity. This study characterizes the microbial and microeukaryote communities, including meiofauna, that inhabit and surround temperate seagrass meadows. By analyzing these communities across different areas of Bodega Harbor, California, we demonstrate how seagrass and environmental factors affect the resilience and overall health of the ecosystem. Our findings highlight the importance of seagrass in maintaining biodiversity and suggest that microscale interactions and habitat variation are important factors shaping these temperate coastal ecosystems. A comprehensive understanding of the variation within benthic communities associated with seagrass habitats can significantly enhance conservation and restoration efforts for seagrass meadows..

## Introduction

Seagrass habitats are distributed worldwide in shallow coastal waters, covering approximately 277,000 km^2^, and span tropical, temperate, and boreal regions (1–3). These highly productive areas support a diverse array of species, from microorganisms to large animals, and are therefore recognized as biodiversity hotspots. Seagrasses offer shelter and food (i.e., nursery grounds) for many economically significant species of fish, crustaceans, and molluscs. They also deliver key ecosystem services, including carbon storage, sediment stabilization and accretion, and coastal protection (1, 2, 4). However, despite their high societal value, seagrasses are declining globally due to various stressors linked to anthropogenic activities and climate change, especially in the intertidal of low-inflow estuaries in temperate regions (4–6).

A variety of abiotic and biotic factors influence biodiversity and community structure in intertidal estuaries, including the presence of seagrasses. Studies have found that fish and benthic invertebrate communities differ between seagrass and bare sediments in both temperate and tropical regions, including changes in organismal density and biomass (7–9). Grazing and bioturbation by larger organisms can also impact seagrass density and spatial distribution, which may, in turn, affect assemblages of smaller-sized benthic species (10, 11). Other abiotic factors, such as those related to daily tide cycles and seasonality (e.g., temperature, salinity, water depth, sunlight exposure), contribute to species richness and spatial patterns of biodiversity by creating environmental gradients (12, 13). At times, these factors result in more pronounced differences among sites along the estuary than between microhabitats (i.e., seagrass vs. bare sediment), highlighting the importance of spatial scale variation (12). In other cases, the differences among sites within an estuary can be just as significant as those observed between microhabitats (14, 15). In other words, abiotic factors can promote larger differences between estuarine sites versus habitat types themselves.

Most research on seagrass habitats has focused on the macrophytes themselves, including factors that control their distribution and abundance, or on their association with other organisms, particularly macro- and megafauna such as fishes, crustaceans, and polychaetes (16). Studies of intertidal seagrass habitats have generally shown that seagrass meadows support higher macroinvertebrate diversity, richness, and density than unvegetated areas (16–18). However, Barnes ((13, 19)) reported little to no effect of seagrass on benthic invertebrates near the mouth of the Kynsa estuary. Notably, differences between seagrass and bare sediment communities increased upstream, influenced by local environmental gradients, which also strongly affect seagrass distribution in estuaries and coastal lagoons (13, 19). Similarly, seagrass meadows often exhibit greater meiofauna abundance and diversity, likely due to higher levels of dissolved oxygen and chlorophyll a (20, 21). Intertidal seagrass habitats have also been shown to increase the stability and richness of microbial communities (22). By creating microhabitats on plant surfaces and within seagrass patches (23, 24), generalist and specialist microbes play key roles in biogeochemical cycles (25–28). Additionally, both emergent and submerged seagrass vegetation are known to vertically structure microbial and macroinvertebrate communities (27, 29).

Understanding the ecological interactions between microorganisms and seagrasses is crucial for effective conservation measures (8, 16, 30, 31). However, a significant gap remains regarding benthic microbes and microbial eukaryotes in intertidal seagrass meadows, resulting in poor understanding of prokaryotic and eukaryotic biodiversity, microhabitat structure, and species co-occurrence in these habitats. Few studies have examined prokaryote and eukaryote co-occurrence in aquatic environments (27, 32–34), and even fewer have addressed these interactions specifically in seagrass habitats (35). While some correlations between prokaryotes and eukaryotes have been observed, the underlying drivers remain unclear. Recent advances in environmental DNA (eDNA) metabarcoding offer a promising approach to filling these knowledge gaps in seagrass ecosystems, particularly in mapping the spatial distribution and interactions of microbial and microeukaryote communities. Studies using multiplexed eDNA metabarcoding—such as 16S ribosomal RNA (rRNA) for bacteria and archaea, 18S rRNA or cytochrome oxidase subunit I (COI) for eukaryotes, and ITS rRNA for fungi—have provided a comprehensive understanding of seagrass biodiversity, facilitating deeper insights into the distribution and relationships among prokaryotic and eukaryotic assemblages (36–39).

In this study, we used 16S rRNA and 18S rRNA eDNA metabarcoding to characterize the microbial and microeukaryote communities associated with seagrass and adjacent bare sediments in Bodega Harbor, Northern California, USA. Through replicated sampling across known environmental gradients in this temperate seagrass habitat, we assessed whether community patterns for microbes and microeukaryotes reflected differences among sites across the estuary or specific microhabitats within sites. We also evaluated the relationships between microbial and microeukaryote communities, focusing on significant co-occurrences of bacteria and microbial metazoa (nematodes) to identify potential spatial patterns of symbiosis.

## Results

### Environmental characterization of Bodega Harbor

We assessed the environmental variables of seagrass and bare sediment samples at three distinct sites at the Bodega Harbor (**Figure 1**). Analysis of environmental variables grouped samples by site and habitat, and primarily highlighted differences in sediment properties (**Figure 2A**). Mason’s Marina, located in the inner harbor, exhibited higher TOC, Silt%, and Clay%, especially in seagrass habitats. Westside Park (mid harbor) and Campbell Cove (outer harbor) were characterized by higher Sand% (**Table S1**). Differences between seagrass and bare sediment habitats varied by site, with Westside Park showing the most pronounced differences: bare sediments had higher Sand%, while seagrass had higher TOC and Sulfate. PC1 and PC2 together explained 99.06% of the total variation in sediment samples (**Figure 2A**). Significant differences were found for all environmental variables between sites, habitats, and habitats within sites (**Table S1 and S4**).

**Figure 1.**
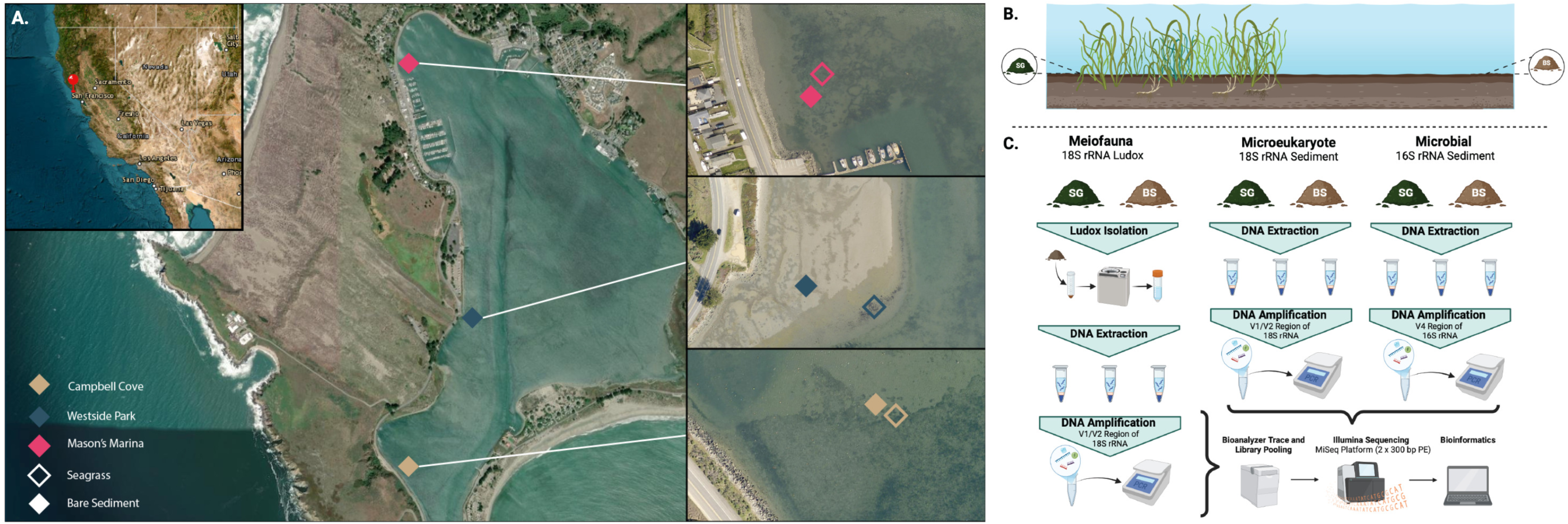
Map and schematic depicting the sampling locations and study design. A) Map depicting sampling sites and microhabitats in Bodega Harbor, California, USA. Pins colors indicate Campbell Cove (tan), Westside Park (blue), and Mason’s Marina (pink). Open symbols represent seagrass, while closed symbols represent bare sediment. B) Schematic depiction seagrass vs bare sediment samples, and C) workflow for the meiofauna, microeukaryotes, and microbial communities. Satellite imagery was obtained from the ArcGIS Map Server via ArcGIS online. Pins for Campbell Cove and Mason’s Marina were offset for clarity due to their close proximity. GPS coordinates for all sampling sites are provided in **Table S1**.

**Figure 2:**
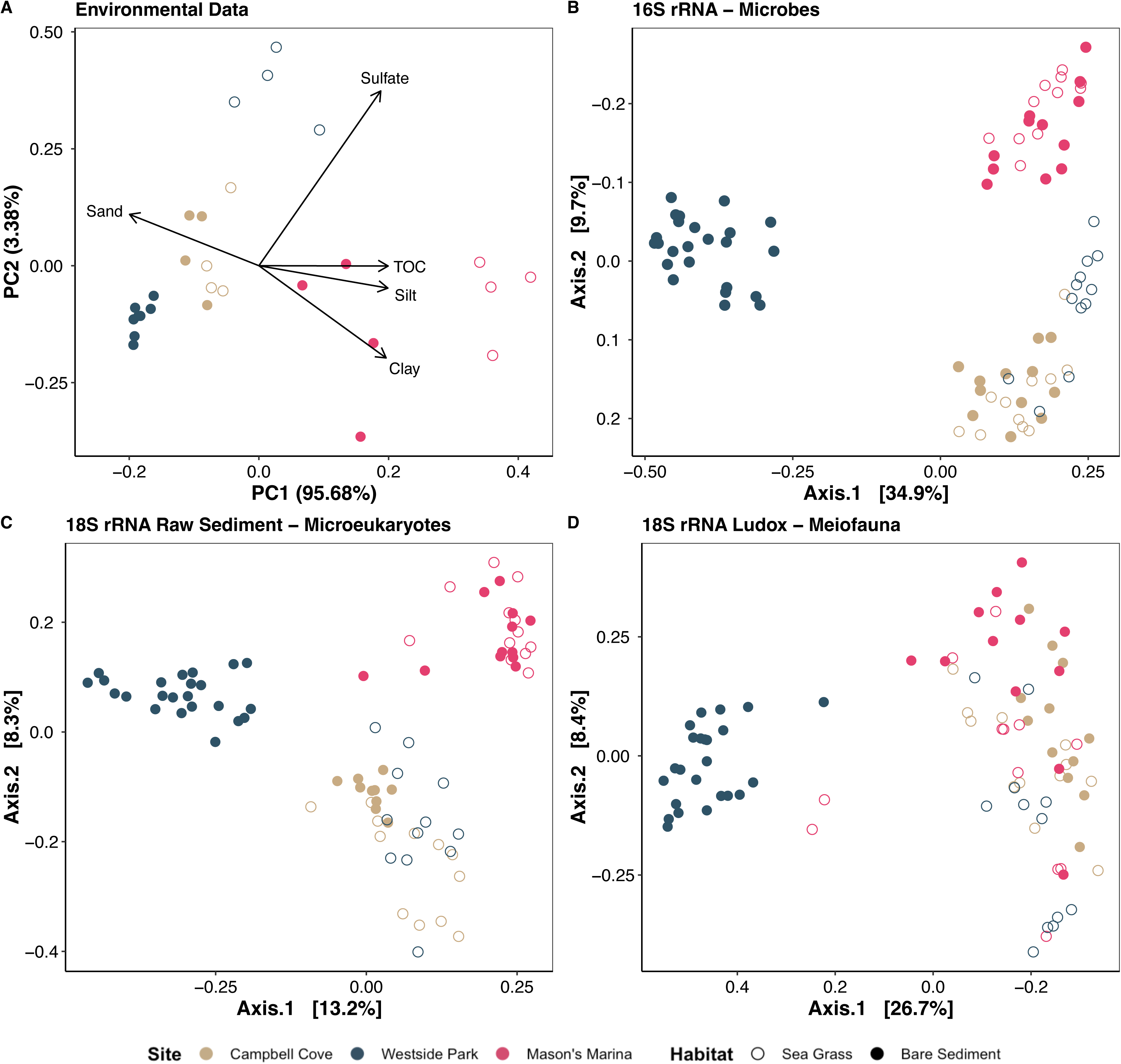
Spatial patterns across datasets. **A**) Principal components analysis (PCA) of environmental data (Euclidean distance from normalized variables). **B-D**) Principal Coordinates Analysis (PCoAs) of (**B**) microbial community (16S), (**C**) broad microeukaryote community (18S raw sediment), and (**D**) meiofaunal community (18S Ludox). PCoAs used Bray-Curtis distance based on ASV transformed data (i.e., relative abundance). Sample sites: Campbell Cove (Tan), Westside Park (Dark Blue), and Mason’s Marina (Pink). Habitats: bare sediment (closed circles) and sea grass (open circles).

### Patterns of alpha- and beta-diversity in microbial and microeukaryotic communities at Bodega Harbor

We compared alpha diversity between bare sediment and seagrass habitats within sites (**Figure S1**). For microbes (16S), there were no significant differences for Observed, Shannon, and Simpson diversity estimates between the two habitats (**Figure S1A**). Westside Park displayed significantly higher evenness in seagrass, while Mason’s Marina had significantly higher evenness in bare sediments (**Figure S1A)**. For microeukaryotes (18S raw sediment), bare sediments had significantly higher Observed diversity compared to seagrass at Campbell Cove and Westside Park (**Figure S1B**). Additionally, bare sediments at Campbell Cove had significantly higher Shannon and Simpson diversity than seagrass (**Figure S1B**). For the meiofauna dataset (18S Ludox), bare sediments had higher Observed diversity compared to seagrass in all three sites (**Figure S1C**). At Westside Park, seagrass had significantly higher Shannon, Simpson, and evenness compared to bare sediments (**Figure S1C**), while at Mason’s Marina, seagrass had significantly lower Shannon and Simpson diversity (**Figure S1C**). Overall, alpha diversity was consistently higher in Campbell Cove than in Mason’s Marina across all datasets, except for Observed and Shannon in the 18S raw sediment dataset (**Figure S6 and Table S3**). Westside Park showed highly variable values, although it had the lowest means for all metrics in the meiofauna dataset. When comparing habitats, bare sediments had higher Observed and Shannon diversity in every dataset. Furthermore, for microeukaryotes (18S raw sediment), bare sediment exhibited significantly higher Simpson (**Table S3)**.

Microbial communities (16S dataset) showed a strong separation by sampling sites, suggesting an environmental gradient from outer to inner sites at Bodega Harbor (**Figure 2B**). Microeukaryotes (18S raw sediment) were also strongly structured by sampling sites, with slightly more overlap between Campbell Cove and Westside Park samples (**Figure 2C**). Meiofaunal communities (18S Ludox) showed a similar pattern but with much more overlap among sample sites (**Figure 2D**). Regardless of the dataset, bare sediment communities from Westside Park strongly differed from all other samples. PERMANOVA analysis and pair-wise tests detected significant differences among sites and between habitats for all datasets (**Table S4**).

### Taxonomic Composition and Differential Abundance of the Microbial Communities and Predicted Gene Functions (16S rRNA dataset)

Microbial communities were predominantly composed of the order *Desulfobulbales*, especially at Campbell Cove (**Figure 3A**). *Desulfobacterales* and *Campylobacterales* were also widespread, except at Westside Park bare sediments where *Pirellulales*, *Rhodobacterales*, and *Rhizobiales* were more prevalent. Among bacterial families, *Desulfocapsaceae* and *Desulfosarcinaceae* dominated across sites, though *Desulfosarcinaceae* was less common in Westside Park bare sediments (**Figure S2A**). *Sulfurovaceae* was the most abundant in Westside Park seagrass, but nearly absent in Westside Park bare sediments. *Pirellulaceae* and *Rhodobacteraceae* were more abundant in Westside Park bare sediments, whereas *Halieaceae* was slightly more prevalent at Mason’s Marina.

**Figure 3:**
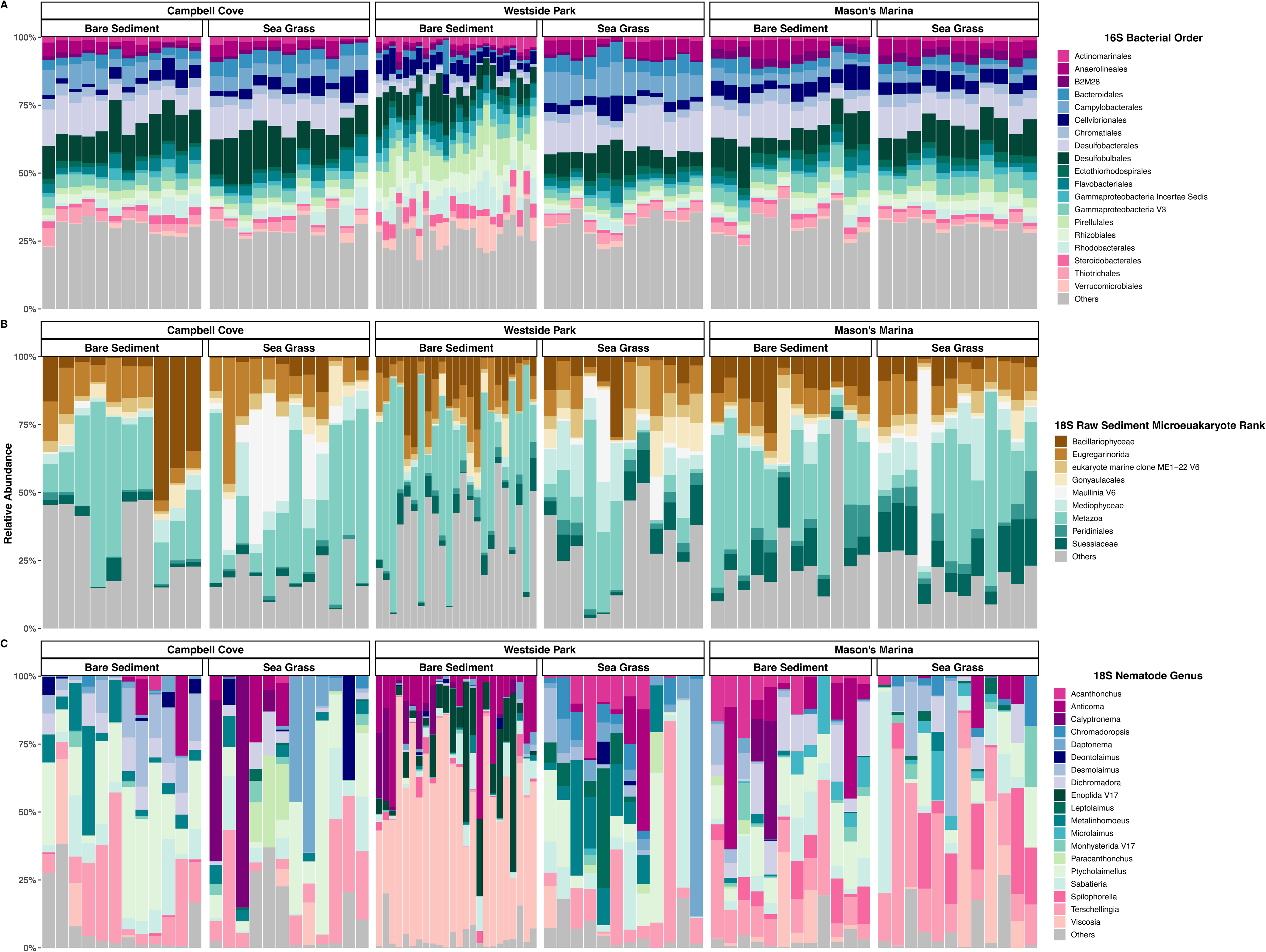
Composition of microbial, microeukaryotic, and nematode communities across sites and habitats in Bodega Harbor. **A**) Order-level classification of the top 19 most abundant bacteria and archaea (16S). **B**) The 9 most abundant microeukaryotic taxa (18S raw sediment). **C**) The 19 most abundant nematode genera recovered from meiofaunal samples (18S Ludox). Lower abundance taxa in each dataset are grouped under “Others”.

Differential abundance analysis detected differences among bacterial genera between sites and among habitats. *Desulfocapsa* and group MSBL-7 (*Deltaproteobacteria*) were enriched at Campbell Cove, while *Sulfurimonas*, *Desulfosarcina*, and *Desulfobulbus* were depleted in Westside Park (**Figure S3**). Westside Park, especially bare sediments, showed enrichment for *Anderseniella*, *Ahrensia*, *Eudoraea*, *Microbulbifer*, and *Parasphingopyxis*. *Halochromatium* was especially enriched at Mason’s Marina. Differentially abundant bacterial families provided insight into nutrient cycling across sites and habitats (**Figure 4**). Those involved in nitrogen (e.g., *Nitrospiraceae* and *Nitrosococcaceae*) and methane cycling were enriched in Westside Park bare sediments, as were *Methylophagaceae* at Campbell Cove and *Methylomonadaceae* at Mason’s Marina. Sulfur cycling families (e.g., *Sulfurimonadaceae* and *Desulfosarcinaceae*) were depleted in Westside Park bare sediments, whereas *Ectothiorhodospiraceae* and *Chromatiaceae* were enriched in seagrass habitats at Westside Park and Mason’s Marina.

**Figure 4:**
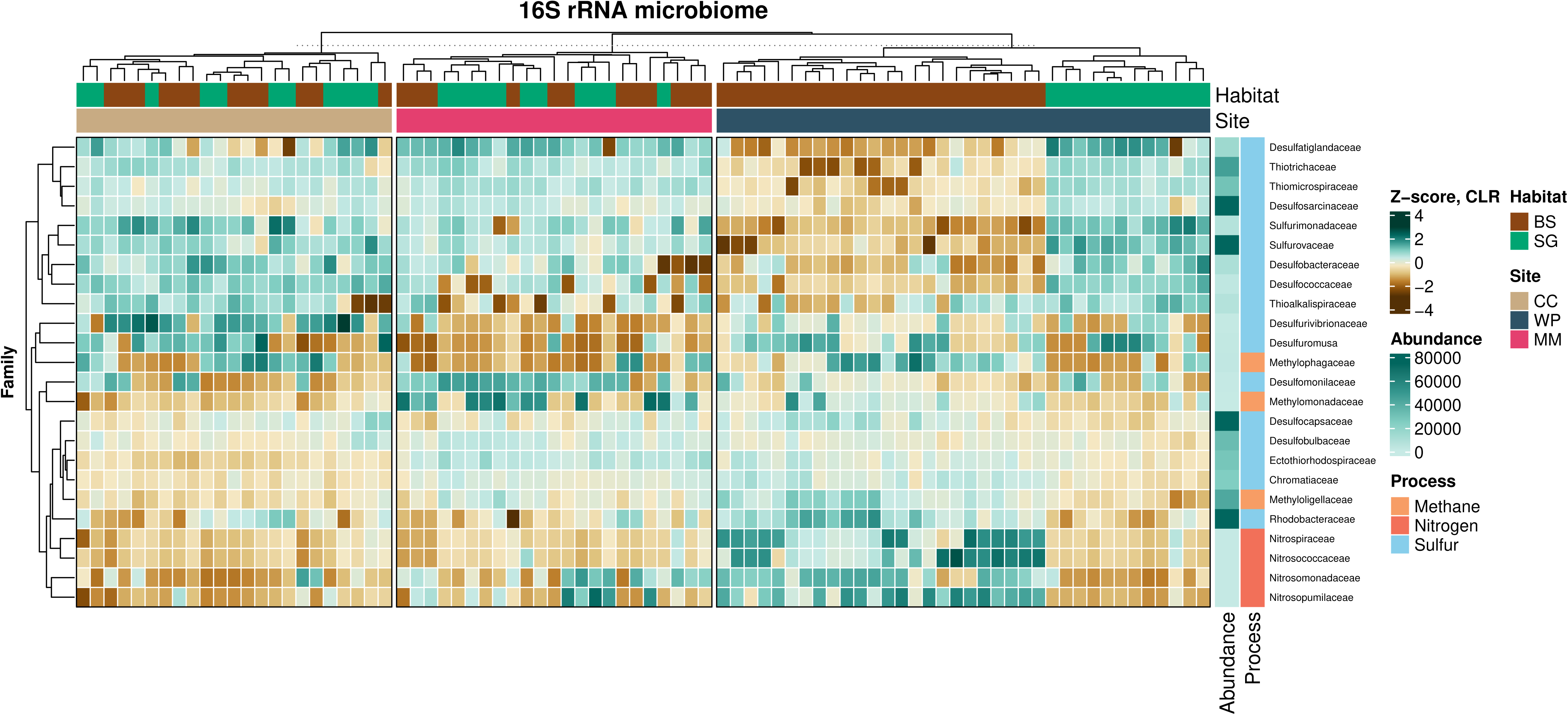
Patterns of differential abundance in bacterial and archaeal taxa involved in biogeochemical cycling. The heatmap shows microbial families associated with methane, nitrogen, and sulfur cycling across sample sites (CC: Campbell Cove, WP: Westside Park, MM: Mason’s Marina) and habitats (BS: bare sediment, SG: seagrass). Family abundance was transformed using the centred-log ratio (CLR). Dark green indicates high abundance, while brown represents low abundance. A complete list of differentially abundant genera is provided in Figure S3.

Overall, there were 1,618 differentially abundant predicted functions among sites, 64 related to sulfur cycling (**Table S5**). Pathways for Dissimilatory Sulfate Reduction (e.g., Dissimilatory sulfite reductase, Adenylyl-sulfate reductase, and Thiosulfate Reductase) were enriched across samples, except at Westside Park bare sediments (**Figure S4**). In contrast, pathways for glycosaminoglycan degradation (e.g., Iduronate−2−sulfatase and N−acetylgalactosamine−4−sulfatase) were more abundant in Westside Park bare sediments. Dimethyl sulfide:cytochrome c2 reductase, a key enzyme in sulfur metabolism, was consistently elevated in Westside Park bare sediments and also enriched at Mason’s Marina. Select sulfur-associated predicted functions were used to analyze patterns in sulfur-utilizing, host-associated bacterial genera (**Table S5**). Westside Park bare sediments had the lowest abundances of sulfur-utilizing host-associated taxa, but the highest *Candidatus Thiobios* (an ectosymbiont of protists; (40, 41)) abundance (**Figure 5A and Table S5**). Sequences matching other potential endosymbionts and *Thiotrichaceae* dominated other sites and habitats.

**Figure 5:**
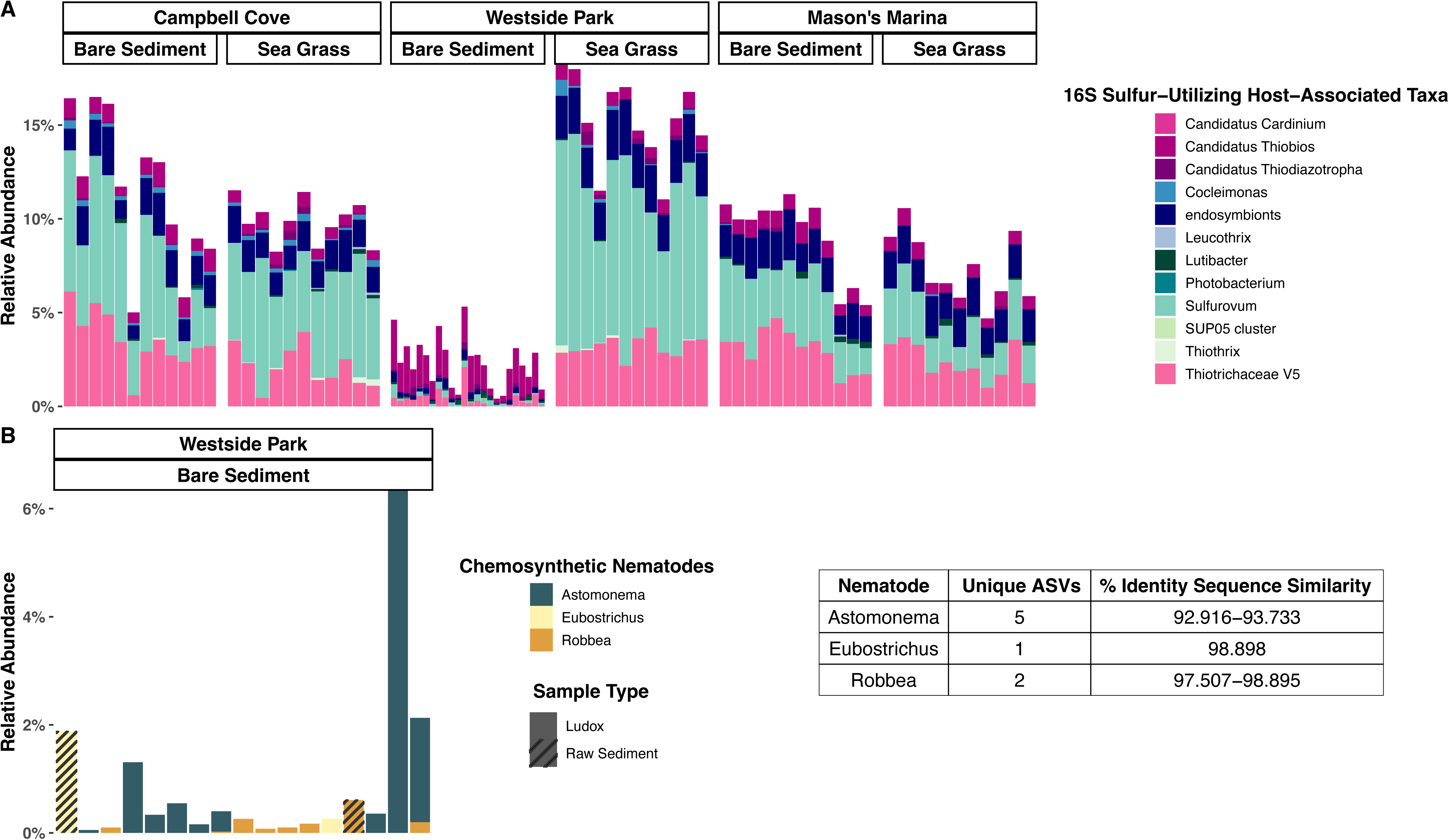
Community composition of chemosynthetic nematode taxa (*Astomonema*, *Eubostrichus*, and *Robbea*) and sulfur-utilizing, host-associated bacteria identified in eDNA datasets. Abundance bar charts show: **A**) host associated bacterial genera known to contribute to sulfur pathways, and **B**) nematode genera within the subfamily Stilbonematinae. Bacterial genera were selected based on their involvement in sulfur pathways of interest (Figure S4 and Table S5), as well as literature describing symbiotic lifestyles. BLAST analysis of “Endosymbiont” sequences identified them as bacterium symbionts of *Alviniconcha*, *Thyasira*, and *Maorithyas hadalis* (all Mollusca). Nematode genera were extracted from both Ludox (15 samples) and raw sediment (2 samples) datasets. For each nematode genus, unique ASV count and percent identity were obtained from DADA2 outputs.

### Major Microeukaryote (18S rRNA Raw Sediment) and Meiofauna Taxa (18S rRNA Ludox)

The 18S rRNA raw sediment dataset was dominated by Metazoa, though other eukaryote groups were also abundant (**Figure 3B**). The parasitic protist order Eugregarinorida was notably more abundant in raw sediment samples compared to the Ludox dataset. Bacillariophyceae diatoms were also commonly found in raw sediment samples, especially in bare sediment habitats. The dinoflagellate group Suessiaceae was particularly abundant in the inner estuary at Mason’s Marina, while the Phytomyxea parasite *Maullinia* was more common in seagrass habitats, especially at Campbell Cove.

Differential abundance analysis detected differences among microeukaryotes between sites and among habitats. Enriched taxa included *Gomphonema rosenstockianum* (diatom) and *Pirsonia formosa* (Stramenopiles) at Campbell Cove, dinoflagellate species *Bysmatrum subsalsum* and *Biecheleria brevisulcata* at Mason’s Marina, *Lecudina* (Alveolates), *Oblongichytrium sp.* (Stramenopiles), and DSGM−81 at Westside Park bare sediments, and *Minidiscus* sp. (diatom) at Westside Park seagrasses (**Figure S5A)**. Metazoa also dominated the 18S Ludox reads (**Figure S2B**), followed by dinoflagellate Gonyaulacales, which was widespread except at Westside Park. Fungi Dikarya were found primarily in seagrass habitats across sites, and the dinoflagellate Peridiniales was most abundant at Campbell Cove.

### Taxonomic Composition of Nematode Communities

A total of 772 ASVs assigned to the phylum Nematoda were recovered from the 18S Ludox dataset, representing 92 genera and 44 families. The most abundant families were Linhomoeidae, Chromadoridae, and Oncholaimidae (**Figure S2C**). Although Anticomidae and Enchelidiidae were commonly detected, they were generally less abundant overall. Oncholaimidae, along with other putative Enoplida taxa that could not be assigned to lower taxonomic levels, were primarily present in bare sediments at Westside Park. Among genera, the most abundant were *Ptycholaimellus*, *Terschellingia*, and *Viscosia* (**Figure 3C**). While *Anticoma* was frequently detected across sites, it was comparatively less abundant. *Viscosia, Calyptronema,* and Enoplida were primarily abundant in bare sediments at Westside Park. *Daptonema* and *Metalinhomoeus* were more prevalent in seagrass habitats at Campbell Cove and Westside Park.

Differential abundance analysis detected differences among nematode genera between sites and among habitats. Enriched taxa at Campbell Cove included *Achromadora, Metalinhomoeus, Terschellingia,* and *Ptycholaimellus*, with the latter two also enriched in Mason’s Marina (**Figure S5B**). Genera enriched at Westside Park included *Anticoma*, Enoplida, and *Viscosia,* especially in bare sediments. Additionally, *Acanthonchus*, *Paramonhystera*, and *Daptonema* were especially enriched in Westside Park seagrass. At Mason’s Marina, unknown Monhysteridae, *Microlaimus,* and *Spilophorella* were more common. Seven unique ASV sequences identified as nematodes known to possess chemosynthetic bacterial symbionts (i.e., *Astomonema*, *Eubostrichus*, and *Robbea*) were also detected in Westside Park bare sediments (**Figure 5B**).

### Co-Occurrence Network Between Bacteria/Archaea and Nematodes

The co-occurrence network, built using strong (r ≥ |0.4|) and significant (p-value < 0.05) correlations between microbial and nematode genera, is composed of 230 nodes and 768 edges (**Table S6**). We focused our analysis on positive correlations to explore potential synergistic relationships between specific microbes and nematodes. This subnetwork contained 203 nodes and 434 edges, with 31 nodes representing nematode genera and 172 nodes corresponding to bacteria/archaea orders (**Figure 6**). The network’s average degree was 4.276 with a modularity of 0.462, indicating a moderate to fairly high level of modular structure. This suggests the presence of well-defined groups of co-occurring bacterial and nematode genera. Supporting this, a large hub in the network included multiple nematode taxa such as Enoplida, *Viscosia*, and *Astomonema*, with smaller hubs made up of one or two nematode taxa. These patterns may reflect microhabitat structures, selective associations, or environmental gradients. Overall, we identified 99 unique bacterial/archaea orders with *Flavobacteriales* (33), *Rhizobiales* (21), *Cellvirbionales* (18), *Bacteroidales* (17), and *Desulfobacterales* (17) showing the highest degree values. Among nematode nodes, Enoplida (92) had the greatest number of connections, followed by *Viscosia* (64), *Astomonema* (37), *Terschellingia* (36), and *Ptycholaimellus* (26). Correlation coefficients for the network ranged from 0.4 (*Calomirolaimus* - d142 V3 [*Acidobacteriota*]) to 0.95 (Enoplida - *Flavobacteriales*).

**Figure 6:**
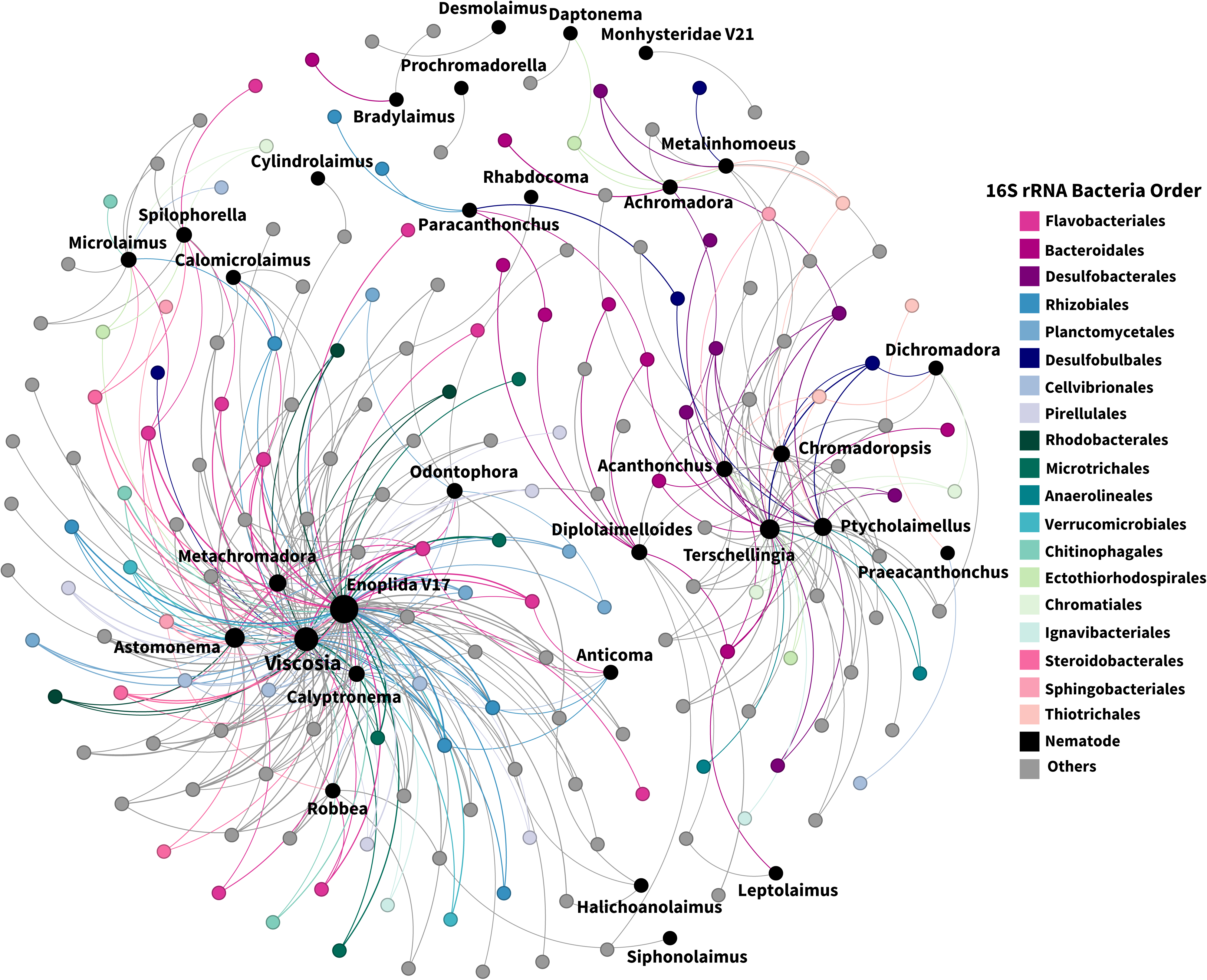
Correlation-base network analysis illustrating potential interactions between bacteria/archaea (16S, order level) and nematodes (18S Ludox, genus level). Edges (lines) between nodes indicate positive co-occurrence relationships (Spearman’s correlation coefficient (ρ) > 0.4; P < 0.05). Node colors distinguish nematodes (black) from various bacteria/archaea orders; orders with fewer than three nodes are grouped as “Others”. Node size indicates the number of connections (degree) for each taxon.

## Discussion

### Benthic Environment Determines Community Assemblages

Our study reveals a pronounced environmental gradient across Bodega Harbor, evidenced by differences in sediment composition and the grouping of benthic communities by site (outer: Campbell Cove; mid: Westside Park; inner: Mason’s Marina) rather than by habitat. Environmental analyses revealed a dominance of coarse sediment (Sand%) at the mouth (Campbell Cove) and mid-estuary (Westside Park), while the inner region contained significantly higher fractions of fine sediments (Silt% and Clay%), and TOC (**Table S1**). This aligns with established estuarine dynamics: high-energy conditions near the mouth retain coarser particles, while finer sediments are transported and deposited upstream (42). Consistently, higher TOC concentrations are found further from estuary mouths (8, 43), and are associated with higher organic matter and reduced water flow at Mason’s Marina (44). Environmental variables also indicated habitat variation, with seagrass areas exhibiting higher fine sediment content, sulfate, and TOC (**Figure 2A and Table S1**), likely due to lowered water velocity that limits sediment resuspension (45). Additionally, the detritus from seagrass enhances carbon storage and influences sulfate reduction rates, raising TOC and sulfate levels (31, 46, 47).

Spatial patterns for microbial, microeukaryote, and meiofaunal communities reinforce the role of site-driven environmental filtering. Microbial community composition shifted from sandy, lower-TOC sites (Campbell Cove and Westside Park) to the fine-grained, organic matter-rich sediments at Mason’s Marina, likely reflecting adaptations to nutrient and oxygen gradients (48), as well as community assembly based on sediment grain size (49, 50) (**Figure 2B)**. Microeukaryotes showed a similar trend, though there was more overlap between Campbell Cove and Westside Park seagrass samples (**Figure 2C**). This overlap likely reflects vegetation influence (51, 52), but overall, microeukaryote communities were shaped more strongly by site differences, which aligns with prior research supporting greater similarity between habitats within the same site (8, 53). The isolation of Mason’s Marina in ordinations suggests stronger environmental filtering, likely influenced by higher TOC, finer sediments, and warmer waters from tidal cycles (44, 54).

Meiofauna communities showed substantial overlap among sites, but were affected by seagrass presence (**Figure 2D**). Within sites, meiofauna differed significantly in alpha-diversity between habitats; although seagrass effects varied between mid (Westside Park) and inner (Mason’s Marina) estuary sites. This variation suggests that abiotic factors may not outweigh vegetation effects for meiofauna (**Figure S1 and Table S3**). Environmental filtering is a key force for larger-body meiofauna such as nematodes (55, 56). Differences in meiofauna and nematode communities by habitat have been observed at multiple spatial scales and are linked to increased organic matter and sediment stability from seagrass (9, 57, 58). Thus, both site and habitat contribute to benthic community structure, but the influence of abiotic factors like salinity and sediment size will vary by benthic biological component (12–14, 59).

Additionally, we found that seagrass only affected evenness at Westside Park and Mason’s Marina, not other alpha-diversity metrics, for microbial communities (**Figure S1A**), contrasting with previous studies showing vegetation presence is a key diversity driver (8, 22, 60). Our findings indicate microbial communities are primarily influenced by location within Bodega Harbor, suggesting that microbes preferentially respond to broader estuarine patterns over microhabitat variation. Microeukaryotes and meiofauna typically showed lower diversity in seagrass compared to bare sediments, although this was not always significant (**Figure S1B-C)**. Notably, Westside Park seagrasses—featuring higher sulfate and finer sediments—exhbited higher meiofauna diversity except for measures of richness. In estuaries, habitat effects are strongest where environmental stress is highest, typically upstream. For example higher macrofauna community diversity in the Knysna estuary shifted from bare sediments near the mouth to seagrass upstream (13). In Bodega Harbor, microeukaryote and meiofauna communities at near-mouth seagrass (Campbell Cove) had lower diversity than bare sediment samples; however, the difference in diversity between bare sediments and seagrass becomes less pronounced towards the inner estuary (Mason’s Marina) (**Figure S1B-C and Table S3**).

### Environmental Filtering and Community Structure in Intertidal Mudflats

In this study, “bare sediments” at Westside Park refer to low and high mudflat samples because no clear unvegetated sediments were visible next to the seagrass habitat during sample collection. Our results highlight the unique features of mudflat habitats in estuaries. Westside Park bare sediment samples had the highest Sand%, and the lowest Clay%, Silt%, sulfate, and TOC, closely resembling Campbell Cove, but markedly different from Westside Park seagrass samples (**Figure 2A and Table S1**). At Westside Park bare sediments, the abiotic factors associated with tidal cycles (e.g, temperature, salinity, sediment moisture) likely drive community differences, rather than the measured sediment properties, which align with their tidal positioning and exposure to hydrodynamics (61). Mudflats display unique zonation patterns due to elevation and tidal exposure; for instance, higher elevated mudflats exposed to greater wave energy tend to retain more coarse sand and less fine sediment (62, 63). Bare sediments at Campbell Cove and Mason’s Marina, sampled adjacent to seagrass beds, may benefit from seagrass-related sediment stabilization and fine particle trapping (45).

Harsh abiotic conditions can significantly affect benthic community composition, primary production, and seagrass biomass (64–67). While shifts in macrofauna taxa do not greatly influence mudflat ecological functioning (68), changes in microbial assemblages can impact metabolic functions (69, 70). For example, enrichment of glycosaminoglycan degradation pathways in Westside Park bare sediments is attributed to greater abundances of the bacteria *Rhodopirellula* and indicates enhanced metabolic capacity for processing marine organic matter (71, 72) (**Figures S3 and S4**). Furthermore, the enrichment of nitrogen cycling bacteria (*Nitrospiraceae*, *Nitrosococcaceae*, *Nitrosomonadaceae*, and *Nitrosopumilaceae*) and to a lesser extent, methane cycling bacteria (*Methylophagaceae*, *Methylomonadaceae*, and *Methylococcaceae*) in Westside Park bare sediments reflect the habitat’s biogeochemical conditions (**Figure 4**). *Nitrospira* species, abundant in mudflat and other marine environments, perform nitrite oxidation under the typical irregular oxic-anoxic conditions in marine sediments (69, 70, 73). Marine mudflats are recognized as habitats for methylotrophic methanogenesis, where methane is produced from methylated compounds such as methanol and methylamines (74–76), suggesting that abiotic conditions of Westside Park bare sediments can facilitate methanogenesis during anoxic periods (77). The low sulfur cycling signal detected in Westside Park bare sediments further underscores differences in organic matter availability and abiotic conditions. In contrast, seagrass beds are known for high sulfur cycling due to oxygen production and increased organic matter (22, 78). Key bacterial taxa, such as *Desulfobacteraceae*, *Thiothrichaeceae*, and *Desulfatiglandaceae* were enriched in all seagrass habitats and bare sediments at Campbell Cove and Mason’s Marina. This finding supports that both seagrass beds and seagrass-adjacent sediments host diverse sulfur-oxidising and sulfate-reducing communities that contribute to sulfide detoxification and nutrient cycling (22, 79).

Meiofauna diversity and community composition were most distinct at the Westside Park site. Although bare sediments had significantly higher observed ASVs, other alpha-diversity metrics were significantly greater in seagrass habitats (**Figure S1C**). Meiofaunal composition in Westside Park mudflats was dominated by Enoplid nematodes including the genus *Viscosia*, reflecting stronger habitat-specific environmental filtering and decreased diversity (80) (**Figure 3C**). The Enoplida order is highly diverse among marine nematodes and includes the largest-bodied taxa, many of which are predators or scavengers (81, 82). *Viscosia* and other enoplids (e.g., *Enoplolaimus litoralis*) are commonly found in sandy intertidal sediments (83–85). While these nematodes typically inhabit the upper sediment layers, they may move deeper when bioturbation occurs or in search of prey (80, 86, 87). Their swimming behavior enables rapid movement within sediments, alternating between active and inactive, which facilitates dispersal through tidal currents (88). Thus, the environmental conditions in Westside Park bare sediments—characterized by larger interstitial spaces and low organic matter—may favor larger predatory nematodes over deposit feeding types (89, 90).

### Symbiotic Sulfur Cycling: Nematode-Bacteria Association Across Estuarine Sites

Seagrass beds are hotspots for sulfur cycling, fostering sulfur-utilizing bacteria and creating conditions for symbiotic relationships (25, 39, 91). Seagrass roots promote these bacteria by generating redox gradients through root oxygen loss and enhancing organic compound availability in sediments (25, 92). In Bodega Harbor, Dissimilatory Sulfate Reduction (DSR) pathways—including Thiosulfate Reductase, Adenylyl−sulfate reductase, and Dissimilatory sulfite reductase— were enriched in seagrass habitats and adjacent bare sediments, except for Westside Park bare sediments (**Figure S4**). The DSR pathway, which can oxidize water-insoluble sulfur particles, is a key function among symbiotic sulfur-oxidizing bacteria (93). We identified particular microbial taxa contributing to these cycles as being host-associtated (**Figure 5A**). For instance, *Candidatus Thiobios* (ectosymbiont of *Zoothamnium niveum*) and *Candidatus Thiodiazotropha* (symbiont of *Loripes lucinalis*) were common in DSR pathways (**Table S5**) (40, 41). Additionally, taxa from the SUP05 cluster, including *Thiotrichaceae* (e.g., *Thiothrix*, *Cocleimonas*, and *Leucothrix*), *Sulfurovum* spp., and unclassified endosymbionts of sea snail *Alviniconcha* and thyasirid clams *Thyasira* and *Maorithyas hadalis* were strong DSR contributors (94–98).

However, chemosynthetic nematodes (i.e., *Astomonema*, *Eubostrichus*, and *Robbea)* associated with the *Thiosymbion* genus were detected only in Westside Park bare sediments (**Figure 5B**). Oncholaimid nematodes, known for hosting diverse symbiotic microbes (99–101), were also enriched there. *Astomonema* are mouthless nematodes harboring *Thiosymbion* in their body cavity (102), relying on these symbionts for nutrition (102, 103). Eubostrichus and Robbea, Stilbonematinae subfamily members, are covered with ectosymbiont *Ca. Thiosymbion bacteria* (104). These sulfur-fixing bacteria, thought to be a food source for nematodes, benefit from nematode movement through sulfide gradients in marine sediments (105–107). The sandy interstitial spaces and variable oxic conditions of Westside Park bare sediments may provide ideal habitats for these nematodes (105).

*Viscosia* (Oncholaimidae) and *Astomonema* were prominent in our nematode-bacteria co-occurrence network, with 64 and 37 positive connections, respectively (**Figure 6**). *Robbea* had eight significant connections. All three nematodes shared bacterial co-occurrences with *Microbulbifer (Cellvibrionales)* and PAUC26f (*Acidobacteriota)* (**Table S6**). *Microbulbifer*, known for degrading polysaccharides and producing broad-spectrum antibacterial alkanoyl imidazoles, was previously isolated from sponges and sea urchins (108, 109) (110–112). PAUC26f is a sponge-associated bacteria, its symbiotic function and evolutionary adaptations remains unknown (113). Other bacterial orders with ≥ 90% symbiotic genomes (114) such as *Bdellovibrionales*, *Rhizobiales*, *Legionellales*, and *Polyangiales* showed notable connections with *Astomonema*, *Robbea*, and *Viscosia*. *Muricauda* (*Flavobacteriales*) uniquely co-occured with *Viscosia* and other Enoplid nematodes, and helps improve coral and clam resilience to stress via antioxidant carotenoid production (115–117). The co-occurrence of chemosynthetic nematodes and likely symbiotic taxa suggests that harsher conditions (e.g., greater exposure to sunlight, higher temperatures) in intertidal mudflats due to tidal cycling in Bodega Harbor may promote such associations, emphasizing the importance of symbiotic-facilitated resilience.

The high connectivity of sulfur-utilizing bacteria in Campbell Cove and Mason’s Marina seagrasses, their adjacent sediments, and Westside Park seagrasses supports the idea that seagrasses create sulfur-rich microenvironments favorable sulfur-based symbioses. Interestingly, the nematode *Diplolaimelloides* was found only in seagrass habitats (**Figure 3C**) and co-occurred with *Campylobacterales* (e.g., sulfur-oxidizing genera *Sulfurimonas* and *Sulfurovum*) and *Oceanospirillales* (aerobic *Marinomonas*). *Campylobacterales* symbionts are known to possess complete sulfur-oxidizing complexes and can fix carbon dioxide via the reverse tricarboxylic acid (rTCA) and Calvin-Benson-Bassham (CBB) pathways using electrons from reduced sulfur (100, 118), also acquiring metabolites like vitamin B_6_ and molybdenum cofactor from hosts (118–121). *Oceanospirillales* have symbiotic relationships with sea urchins, corals, mussels, and bone-eating worms, providing hydrocarbon degradation (122, 123). *Diplolaimelloides–Oceanospirillales* associations may support seagrass resilience and carbon sequestration (1, 2, 4, 124). Sulfur-oxidizing *Sulfurimonas* and *Sulfurovum* are known to provide carbohydrates and amino acids to host invertebrates or bacterivores, aiding environmental adaptation (97, 125–128).

### Nematode Bioindicators of Environmental Stress and Sediment Quality

Nematodes have been widely used as bioindicators to monitor environmental changes in various aquatic systems, as they can quickly respond to chemical and physical environmental stressors (129, 130), including pollution to heavy metals (131, 132). High concentrations of organic carbon pollutants in sediments affect nematode density and diversity (54, 90, 133, 134). In Bodega Harbor, the nematode genera *Microlaimus* and *Spilophorella* were highly abundant in Mason’s Marina habitats, but depleted in other sites (**Figure S5B**). Both genera had notable associations with *Ectothiorhodospirales* and *Chromatiales* in our co-occurrance network analysis—including purple sulfur bacteria (PSB) that utilize reduced sulfur compounds for anoxygenic photosynthesis (135, 136) (**Figure 6)**. Uniquely, *Microlaimus* and *Spilphorella* shared a connection with *Thiogranum* (*Ectothiorhodospirales),* a chemolithoautotrophic bacterium often linked to high concentrations of heavy metals and nutrients (137). *Microlaimus*, a commonly used bioindicator, is known to be susceptible to copper, but resilient in environments with hydrocarbons, such as mariculture and harbors (129, 138–140). The co-occurrence of these nematodes and *Thiogranum* suggests that these taxa are adapted to habitats shaped by active sulfur cycling and high organic matter turnover at sites such as Mason’s Marina (126, 138–140). *Terschellingia* and *Ptycholaimellus*, found mainly outside of Westside Park bare sediments, may indicate metal-contaminated environments, and are known for tolerating mariculture and persisting in harbors. Notably, *Terschellingia* is resistant to anthropogenic pollution (90, 129, 141, 142), but remains sensitive to physical disturbance and harsh environments, supporting their lower abundances in Westside Park bare sediments (66, 129). Similar spatial trends were seen in Campbell Cove and Westside Park for *Metalinhomoeus* and *Daptonema*, with *Metalinhomoeus* inhabiting high-organic habitats and *Daptonema* being resistant to anthropogenic disturbances (129, 142–144). *Daptonema* is typically found in environments contaminated with diesel and metals (129, 139, 143); however, these nematodes were largely absent from Mason’s Marina (**Figures 3C and S5B**). This may be due to the increased susceptibility to organic enrichment, as their abundance decreased or they were absent in sites with mean organic content higher than 2.80% (90). Mason’s Marina had the highest organic content in Bodega Harbor, suggesting that *Daptonema* and *Metalinhomoeus* may serve as bioindicators for organic content in estuarine ecosystems. The nematode genus *Achromadora* is often absent from habitats with high Zn, As, Pb, or elevated acidity (145), and decreases in abundance from alkaline to acidic soils (146). In Bodega Harbor, *Achromadora* was only found in Campbell Cove where sulfate levels were the lowest, except for Westside Park bare sediments. Increase of sulfate promotes sulfate reduction and accumulation of reduced sulfur in sediments, contributing to acidification and deoxygenation (147, 148). Our results indicate that *Achromadora* is a bioindicator of low pH, low sulfate, or reduced sulfur metabolic pathways. Similarly, *Dichromadora* was enriched in Campbell Cove, present in Mason’s Marina, and depleted in both Westside Park habitats. This genus is considered resistant to general anthropogenic disturbances, pollution, and organic enrichment (129). Further investigation is needed to fully understand the distribution of *Dichromadora* in Bodega Harbor; its absence from Westside Park may relate to the lower levels of TOC.

## Materials and Methods

### Study area

Bodega Harbor, located in Northern California, USA (38°19’59“N 123°02’53”W), is approximately 105 km north of San Francisco. This small low-inflow estuary (∼ 5 km^2^) features extensive intertidal mudflats and limited fringing salt marshes. Finer sediment and higher organic content are found in the inner part of the harbor, whereas sandy sediment dominates near the harbor mouth (44). The estuary is relatively shallow (∼ 2 m of depth at high tide) and connects to the Pacific Ocean via a dredged channel in the northern part of Bodega Harbor (149). The system is strongly affected by the adjacent ocean, with up to 80% of water exchanging during each tidal cycle (149, 150). Freshwater input from small streams is minimal and occurs mostly during the winter rainy season. Mean summer water temperatures range from 15.9 to 18.1 °C (44, 150).

### Sampling

Sediment samples were collected at low tide from Bodega Harbor on July 7th, 2018 across three different sites: Mason’s Marina (inner estuary), Westside Park (mid estuary), and Campbell Cove (outer estuary). At each site, four sediment cores were taken from vegetated (seagrass) and nonvegetated (bare sediment) habitats using a 10 x 10 cm PVC core (**Figure 1**). Samples were placed in Ziploc bags, kept cool in an ice chest, and transported to the laboratory, where they were stored at -80 °C until processing.Vegetated sediments at Mason’s Marina and Westside Park were covered in eelgrass (*Zostera marina*), while those at Campbell Cove also contained sea lettuce (*Ulva lactuca*). At Westside Park, no clearly unvegetated sediments were found adjacent to the seagrass habitat, so “bare sediments” here refer to both low and high mudflat samples. However, for the analysis, these samples were treated as “bare sediments”. In the laboratory, sediment cores were split in half for molecular procedures and sediment analysis. Sediment characterization measured sand (Sand %), silt (Silt %), and clay (Clay %), total organic carbon (TOC), and sulfate sulfur (SO_4-_S). All sediment analyses were performed at the UC Davis Analytical Laboratory (**Table S1**). Sediment samples for molecular work were processed two weeks after collection.

### Extraction of meiofauna from sediment samples

To characterize meiofauna communities, with an emphasis on marine nematodes, sediment samples were washed using filtered artificial seawater (Instant Ocean, Spectrum Brands, Inc., USA) over a 45 μm sieve. Organisms were extracted from the sieved sediment by flotation using Ludox^TM-50^ [specific gravity of 1.15, (151)] following protocols described in Somerfield and Warwick (152). To minimize cross-contamination, sieves and glassware were sterilized and thoroughly washed between each core. The equipment was washed with Milli-Q ultrapure water, soaked in a 10% Sodium Metabisulfite solution (Pro Supply Outlet, CA, USA) for 1 hr, and rinsed in Milli-Q ultrapure water before processing the next sample (153). The final volume in each falcon tube was adjusted to 20 mL; a 1 mL aliquot of the meiofauna community was set aside for morphological identification and DNA barcoding of nematodes as described in Pereira et al. (154). DNA sequences are available in GenBank under accession numbers PV244184-PV244287 (155). The remaining sample in each Falcon tube was centrifuged at 5000 rpm for 2 min to concentrate the organisms into a single pellet. After centrifugation, the supernatant was carefully discarded, and community DNA was extracted from the entire pellet (see below).

### Metabarcoding of microbial and microeukaryote communities

DNA was extracted from raw sediment (0.25 g of sediment) and Ludox-processed samples in triplicate using the ZymoBIOMICS™ DNA Miniprep Kit (Catalog Nos. D4300, Zymo Research Corp, Irvine, CA) according to the manufacturer’s protocol. To monitor potential kit contamination, blank controls containing water were included during the DNA extraction. Lysates were stored at -80°C until PCR amplification. For microbes, the V4 region of the 16S rRNA gene was amplified with primers 515F (5’-GTGCCAGCMGCCGCGGTAA-3’) and 806R (5’-GGACTACHVGGGTWTCTAAT-3’) (156). For the microeukaryotes (raw sediment and Ludox), a 400-bp segment of the V1–V2 hypervariable regions of the 18S rRNA gene was amplified using primers F04 (5’-GCTTGTCTCAAAGATTAAGCC-3’) and R22 (5’-GCCTGCTGCCTTCCTTGGA-3’) (157). All PCRs were performed in a sterilized laminar-flow hood. PCRs had (25 µl total volume) consisted of 1 µl of DNA template, 0.5 µl of each primer (10 µM), 10 µl of Platinum Hot Start PCR Master Mix (2x) (Thermo Fisher), and 13 µl of molecular-grade water. Positive (ZymoBIOMICS Microbial Community Standard, Zymo Research, Irvine, CA) and negative controls (molecular-grade water) were included. PCR amplification followed the protocols described in Pereira et al. (154). DNA concentrations were quantified with a Qubit 3.0 fluorometer with the Qubit® dsDNA HS (High Sensitivity) Assay Kit (Thermo Fisher Scientific) and normalized before library pooling. Libraries were purified using magnetic beads and size-selected (300–700 bp) on a BluePippen (Sage Science). Sequencing was carried out at the UC Davis Genomics Core on the Illumina MiSeq Platform (2 x 300 bp PE) in two runs. All wet lab protocols and bioinformatics scripts are available on GitHub (https://github.com/BikLab/bodega-bay-eDNA).

### Processing Illumina data using DADA2

Raw Illumina datasets were demultiplexed using a custom script to handle dual-barcode combinations. Primer sequences were trimmed with the *cutadapt* plugin (158). Amplicon sequence variants (ASVs) were denoised and estimated using DADA2 (159) within QIIME2 v2024.2 (160) with default parameters and by truncating forward and reverse reads at 250 bp and 200 bp, respectively. Taxonomy for 16S ASVs was assigned using the BLAST+ consensus taxonomy classifier (161) with default parameters (top 10 hits, minimum confidence of 0.80, and a minimum consensus of 0.51), with the 16S rRNA SILVA 138 SSURef NR99 database as reference (162). For 18S ASVs, sequence reads were truncated at 209 bp (forward) and 230 bp (reverse), and taxonomy was assigned using the BLAST+ top hit approach (161) with a minimum confidence of 0.90 as recommended for meiofaunal communities with limited representation in reference databases (155). A custom reference dataset (https://github.com/BikLab/MeioDB-18SrRNA) was used for 18S ASVs, composed of the SILVA 138 SSURef NR99 database with added short (∼350 bp) and large (∼700 bp) fragments covering the V1-V2 regions of the 18S rRNA retrieved from NCBI, improving the accuracy and precision of nematode taxonomy assignments (155).

### Bioinformatics and statistical analysis

Samples with low read count or suspected errors from DADA2 denoising were excluded from downstream analysis (**Table S2**). Potential contaminants were identified using the R package *decontam* v1.28 (163) with the prevalence method and a contaminant classification threshold of 0.5 (**Table S2**). In addition to blanks and controls, samples “MM.SG.1.3” and “CC.SG.3.1” were removed from the 16S rRNA raw sediment dataset, and “CC.B.1.1.RS”, “CC.B.3.3.RS”, and “WP.MFL.2.1.1.RS” from the 18S rRNA raw sediment dataset due to low read counts and suspected naming errors. For the 16S rRNA data, reads identified as mitochondrial, chloroplast, or unassigned at the domain level were removed. For the 18S rRNA data (Ludox and raw sediment), reads annotated as “humans”, unassigned, Charophyta (seagrass), and *Ulva* (green algae) were excluded to focus on microeukaryotes.

Quality-filtered ASV tables were used to assess the microbial and microeukaryote community compositions across habitats (seagrass and bare sediments) and sites (i.e., Campbell Cove, Westside Park, and Mason’s Marina). Alpha-diversity metrics—observed diversity, Shannon diversity *H’* (Log_2_), Inverted Simpson (*D*), and Pielou’s Evenness (*J’*)—were calculated using *phyloseq* v1.52 (164) and compared among habitats and sites. A Kruskal-Wallis (KW) test, followed by Wilcoxon pairwise analyses, were performed. The 16S rRNA sample “CC.SG.3.2” (Campbell Cove seagrass) habitat was excluded from the alpha diversity analysis due to excessive reads (420,654) compared to the dataset mean (21,256 ± 8,172).

Beta-diversity was assessed using Bray-Curtis dissimilarity and ASV relative abundances, and visualized via Principal Coordinate Analysis (PCoA) with *phyloseq* (164). Permutational analysis of variance (PERMANOVA) was used to test for differences (p-value < 0.05) among sites and habitats. Principal components analysis (PCA) using Euclidean distance matrix based on standardized environmental variables was used to characterize sites and habitats. To identify taxa (16S: bacteria, 18S raw sediment: microeukaryotes, 18S Ludox: nematodes) that contribute most to community differences among sites and between habitats, differential abundance analyses were performed at different taxonomic ranks using the ALDEx2 v1.40 (165) after centered-log ratio (CLR) transformation. Significance was assessed by KW tests (p < 0.05), and false discovery rates (FDRs) were estimated via the Benjamini-Hochberg (BH) procedure (166). Functional profiles of bacterial communities (16S rRNA) were predicted with PICRUSt2 v2.6.2-foss-2023a (167), with sulfur cycling/metabolism pathways analyzed separately as above.

Barplots for 16S and 18S rRNA datasets were generated at multiple taxonomic ranks and less abundant taxa were grouped as “Other”. Known sulfur-utilizing taxa with host associations were further visualized. Meiofauna and nematode diversity were analyzed in greater detail in the Ludox 18S dataset, including BLAST (161) and phylogenetic placement (using SILVA ACT (168, 169)) of chemosynthetic nematode ASVs (*Astomonema*, *Eubostrichus*, *Robbea*). *Astomonema* had five unique ASVs ( 92.916%–93.733% similarity), *Eubostrichus* one ASV (98.898%), and *Robbea* two ASVs (97.507%–98.895%).

To examine co-occurrence between bacteria (16S dataset) and nematode genera (18S Ludox dataset) in 82 shared samples, networks were constructed using Spearman correlation coefficient (*r* ≥ |0.4|, p < 0.05 after FDR-corrected via BH procedure) with the HMSC v.5.2-3 package. A prevalence threshold (≥ 5%) was applied and taxa were agglomerated at the genus level and transformed to relative abundances. Networks were visualized with Gephi v0.10.1 (170) using the Fruchterman–Reingold algorithm (171); nodes represented bacteria (colored, labeled at the order level) and nematodes (labeled by taxon). For this study, all visualizations were created in *ggplot2* v3.5.2 (172) using *R* version 4.5.0 (173).

## Acknowledgments

We greatly thank John J. Stachowicz at UC Davis for his assistance in collecting sediment samples for this study in Bodega Harbor. Funding for this study was provided by the Gordon and Betty Moore Foundation (Symbiosis in Aquatic Systems Initiative, grant #9326), and a National Science Foundation CAREER award (DEB-2144304) to HMB at UGA. Research support for ADS was provided by the University of Georgia Research Foundation and the National Institute of General Medical Sciences of the National Institute of Health under award number 1T32GM142623. We thank the Georgia Advanced Computing Resource Center (GACRC) at UGA (https://gacrc.uga.edu/) for computational resources that have contributed to the results in this publication.

## Data availability

Raw amplicon sequences generated in this study have been deposited in the NCBI Sequence Read Archive (Bioproject: PRJNA1369552). All scripts and files required for the reproducibility of all analyses conducted in this study are available via GitHub (https://github.com/BikLab/bodega-bay-eDNA)

## Supplemental figures

**Figure S1: Variation of alpha diversity metrics between bare sediment and seagrass habitats within sites.** The Observed (i.e., Richness), Simpson, Shannon, and Evenness diversity metrics were estimated for both habitats sampled at each location for **A**) microbial (16S), **B**) microeukaryotes (18S raw sediment), and **C**) meiofaunal (18S Ludox) communities. Only significant differences (*p* < 0.05) between habitats, based on the Wilcoxon signed-rank test, are shown (Table S3).

**Figure S2: Composition of microbial, microeukaryotic, and nematode communities across sites and habitats at Bodega Harbor. A**) Family-level classification for the 19 most abundant bacteria/archaea families (16S). **B**) The 9 most abundant meiofaunal taxa (18S Ludox). **C**) The19 most abundant nematode families identified in meiofaunal samples (18S Ludox). Lower abundance taxa in each dataset were grouped under the category “Others”.

**Figure S3. Differential abundance analysis of bacteria and archaea (16S; genus level).** Heatmap of differential abundant bacterial/archaeal genera across sample sites (CC: Campbell Cove, WP: Westside Park, MM: Mason’s Marina) and habitats (BS: Bare sediment, SG: Seagrass). Analysis was run comparing sample sites. ASV abundances were transformed using the centred-log ratio (CLR). Dark green indicates high abundance; brown indicates low abundance.

**Figure S4. Differential abundance analysis based on predicted functions related to sulfur cycling pathways.** The heatmap shows differentially abundant sulfur cycling pathways across sample sites (CC: Campbell Cove, WP: Westside Park, MM: Mason’s Marina) and habitats (BS: Bare sediment, SG: Sea grass). Differential abundance analysis comparing sample sites was first run on all predicted functions (i.e., EC numbers), after which only significant (kw.ep < 0.05) sulfur specific pathways were selected. The abundance of predicted functions (EC numbers) was transformed using the centred-log ratio (CLR). Dark green indicates high abundance, while brown indicates low abundance. A complete list of results for all metabolic pathways is provided in Table S5.

**Figure S5. Differential abundance analysis of microeukaryotes (18S raw sediment; V9 level) and nematodes (18S Ludox; genus level).** Heatmap of differential abundant **A**) microeukaryotes and **B**) nematode genera across sample sites (CC: Campbell Cove, WP: Westside Park, MM: Mason’s Marina) and habitats (BS: Bare sediment, SG: Seagrass). Analysis was run comparing sample sites. ASV abundances were transformed using the centred-log ratio (CLR). Dark green indicates high abundance; brown indicates low abundance.

**Figure S6: Overall alpha diversity metrics across sites in Bodega Harbor.** The Observed (i.e., Richness), Simpson, Shannon, and Evenness diversity metrics were estimated for **A**) microbial (16S), **B**) broad microeukaryote (18S raw sediment), and **C**) meiofaunal (18S Ludox) communities at three sites in Bodega Harbor (CC: Campbell Cove, WP: Westside Park, MM: Mason’s Marina). Only significant differences (*p* < 0.05) among sites, identified by pairwise analysis following a Kruskal-Wallis test, are indicated.

## Supplemental Tables

**Table S1. Environmental variables associated with sites and habitats sampled at Bodega Harbor.** The variables included sulfate, total organic carbon (TOC), sediment texture (percentages of sand, silt, and clay). GPS coordinates for each site and habitat are provided, as well as disturbance levels (i.e., none-low [NL], low-medium [LM], medium-high [MH]), determined by their distance from anthropogenic activity. Mean values of environmental variables across sites and habitats are included. KW analysis results for comparisons between sites, habitats, and individual habitats within sites are presented (significant values in bold).

**Table S2. Sequencing depth for samples representing sites and habitats sampled at Bodega Harbor. The datasets include 16S (bacteria/archaea), 18S raw sediment (broad microeukaryotes), and 18S Ludox (meiofauna).** Data presented are the number of reads (i.e., input) fed to and retained (non-chimeric) by DADA2 as well as the number of unique DNA sequences (ASVs). Final counts of reads and ASVs (i.e., after *decontam*) were used in statistical and ecological analyses. Samples highlighted in red had low read count or possible errors and were excluded from the respective datasets. Samples highlighted in blue are controls and blanks.

**Table S3. Summary of alpha diversity metrics.** Mean, standard deviation (SD), and KW test results (X^2^ and p-value) are presented for each alpha diversity index (Observed, Shannon, Simpson, and Evenness) across all datasets (16S, 18S raw sediment, and 18S Ludox) are provided. Metrics were compared among sites, between habitats, and between habitats within sites for each dataset. Significant p-values are highlighted in bold.

**Table S4. Summary PERMANOVA results, including pairwise comparisons across datasets (Environmental, 16S, 18S Ludox, and 18S raw sediment).** Abbreviations: df: degrees of freedom; SS: sum of squares; R2: coefficient of determination; F: F statistic; Pr(>F): p-value; Res: residual. Sites:Campbell Cove (CC), Westside Park (WP), and Mason’s Marina (MM). Habitats: Bare Sediment (BS) and Seagrass (SG).

**Table S5. Significant differentially abundant predicted functions (i.e., 16S dataset) among sites (kw.ep < 0.05).** Highlighted functions related to sulfur cycling pathways and were presented in Figure S4. Specific sulfur functions of interest were subsetted and further explored for bacterial taxa that are host-associated or hypothesized to be symbionts.

**Table S6. Complete node and edge table for significant correlations (p-value < 0.05) in the co-occurrence network analysis.** The weight represents the Spearman’s correlation coefficient.

